# Not fleeting but lasting: Limited influence of aging on implicit adaptative motor learning and its short-term retention

**DOI:** 10.1101/2023.08.30.555501

**Authors:** Pauline Hermans, Koen Vandevoorde, Jean-Jacques Orban de Xivry

**Affiliations:** KU Leuven, Department of Movement Sciences, B-3000 Leuven, Belgium; KU Leuven, Leuven Brain Institute, B-3000, Leuven Belgium

## Abstract

In motor adaptation, learning is thought to rely on a combination of several processes. Two of these are implicit learning (incidental updating of the sensory prediction error) and explicit learning (intentional adjustment to reduce target error). The explicit component is thought to be fast adapting, while the implicit one is slow. The dynamic integration of these components can lead to an adaptation rebound, called spontaneous recovery: the trace of a first, longer learned adaptation reappears after it is extinguished by a shorter period of de-adaptation. The slow implicit process is still decaying from the first adaptation, resulting in the before mentioned adaptation rebound. Trewartha et al. (2014) found that older adults show less spontaneous recovery than their younger controls, indicating impairments in implicit learning. This disagrees with evidence suggesting that the implicit component and its retention does not decline with aging.

To clarify this discrepancy, we performed a conceptual replication of that result. Twenty-eight healthy young and 20 healthy older adults learned to adapt to a forcefield perturbation in a paradigm known to elicit spontaneous recovery. Both groups adapted equally well to the perturbation. Implicit adaptation of the older subjects was indistinguishable from their younger counterparts. In addition, our conceptual replication failed to reproduce the result of Trewartha et al. (2014) and found that the spontaneous recovery was also similar across groups. Our results reconcile previous studies by showing that both spontaneous recovery and implicit adaptation are unaffected by aging.

## Introduction

Young healthy adults can adapt to a change in the environment and optimize their reaching performance (Morehead and Orban de Xivry 2021; Shadmehr et al. 2010). Such adaptation process of upper limb movements is studied in the laboratory via perturbation of the visual feedback about the moving direction of the hand (Krakauer et al. 2005; Orban de Xivry and Lefèvre 2015), by shifting the visual field via prism goggles (Welch 1969) or by applying a force on the moving arm (Lackner and DiZio 1994; Shadmehr and Mussa-Ivaldi 1994). For any of these perturbations, young participants can readily decrease the effect of the perturbation on their reaching performance through a combination of explicit strategies and implicit adaptation (Morehead et al. 2015; Taylor et al. 2014; Taylor and Ivry 2011; Welch et al. 1974). Implicit adaptation is the incidental updating of the movement driven by sensory prediction error and occurs gradually (Mazzoni and Krakauer 2006; Morehead et al. 2017; Taylor et al. 2014). Explicit adaptation consists of the application of cognitive strategies to reduce target error and reduces errors rapidly (Morehead and Orban de Xivry 2021). The explicit component contributes more to total adaptation for visuomotor rotation than for force-field adaptation. Learning to counteract a force field is largely an implicit process with only a small explicit component (Schween et al. 2020).

Older adults show lower levels of total motor adaptation than young adults (Aucie et al. 2021; Bakkum et al. 2021; Bock 2005; Cressman et al. 2010; Hegele and Heuer 2010; Malone and Bastian 2015; Sombric and Torres-Oviedo 2021)(Buch et al. 2003; Heuer and Hegele 2008; Li et al. 2021; Seidler 2006, 2007; Vandevoorde and Orban de Xivry 2019). Recent evidence suggests that this impairment in motor adaptation is specific to the explicit component of adaptation (Bock and Girgenrath 2006; Hegele and Heuer 2010, 2013; Heuer and Hegele 2008; Li et al. 2021; Vandevoorde and Orban de Xivry 2019, 2020; Wolpe et al. 2020) and that the implicit component of motor adaptation elicited by a visuomotor adaptation and its short-term retention remains unimpaired up to 60-70 years old (Huang et al. 2017; Reuter et al. 2020; Tsay et al. 2023; Vachon et al. 2020; Vandevoorde and Orban de Xivry 2019).

Few studies have investigated the effect of age on force-field perturbation (Cesqui et al. 2008; Huang and Ahmed 2014; Kitchen and Miall 2021; Reuter et al. 2018; Trewartha et al. 2014). Little difference in the amount of adaptation to a force-field perturbation has been found between young and older participants (Huang and Ahmed 2014; Trewartha et al. 2014). Yet, the explicit and implicit components of adaptation have never been measured in these studies. While we know that the contribution of explicit strategies to force-field adaptation is small (Schween et al. 2020), it is not null. Therefore, it remains unknown whether the implicit component of motor adaptation remains unaffected in older people during a force-field adaptation task. Measuring the explicit and implicit components of motor adaptation is essential in order to gain insight into the source of possible deficits.

Interestingly, one study reported a very specific age-related impairment in force-field adaptation. That is, while initial adaptation was unimpaired with age, its short-term retention as measured by spontaneous recovery of adaptation was impaired (Trewartha et al. 2014).

Spontaneous recovery occurs when motor adaptation to some perturbation A, which is then hidden from view due to adaptation of a second perturbation B, reappear without any additional exposure to perturbation A (Coltman et al. 2019; Ethier et al. 2008; Kojima et al. 2004; McDougle et al. 2015; Sarwary et al. 2018; Smith et al. 2006). It suggests that the motor memory of the adaptation to perturbation A is not washed out by adaptation to perturbation B but is retained. Therefore, such spontaneous recovery of motor memories linked to perturbation A represents a proxy for short-term retention of the associated motor memory (Smith et al. 2006).

The presence of spontaneous recovery indicates the presence of at least two learning processes working on different time scales. One process learns and forgets quickly, while the other is slow (Kording et al. 2007; Smith et al. 2006). In this framework, the spontaneous recovery of motor memory of field A is attributed to the memory trace of the slow learning process, (McDougle et al. 2015; Smith et al. 2006). This memory trace is masked by the fast adaptation process during the deadaptation period. Interestingly, the slow process has been associated with the implicit component of adaptation while the fast process has been linked to the explicit component (McDougle et al. 2015).

Three major concepts reviewed up to here bear some contradictions: 1) implicit adaptation and its short-term retention are not impaired by aging (Vandevoorde and Orban de Xivry 2019), 2) spontaneous recovery is impaired in older people (Trewartha et al. 2014), and 3) the slow process of adaptation, which determines spontaneous recovery, corresponds to the implicit component of adaptation (McDougle et al. 2015). In other words, if implicit adaptation is unimpaired in older people and if it determines spontaneous recovery, then spontaneous recovery cannot be different across age groups. Yet, it is unclear where the contradiction comes from as these different studies have marked differences in protocol, which could affect the results. Implicit adaptation level was obtained using a visuomotor rotation of the cursor feedback (Vandevoorde and Orban de Xivry 2019), whereas its short-term retention was measured in a force-field paradigm (Trewartha et al. 2014). As it is known that perturbation type influences implicit adaptation level (Morehead et al. 2015; Schween et al. 2020), this difference in perturbation type could be responsible for this discrepancy. There is thus a need to test these three observations within a single experiment. Therefore, we set out to measure both implicit adaptation and its retention via spontaneous recovery in a single force field paradigm in both healthy young and older adults in order to test four different hypotheses: 1) implicit adaptation levels at the end of the adaptation period are similar across age groups; 2) spontaneous recovery is larger in young than in older participants, 3) implicit adaptation at the end of the adaptation period is correlated with the amount of spontaneous recovery and, 4), as suggested by Trewartha and colleagues, spontaneous recovery is related to explicit memory processes such as working memory capacity.

## Methods

### Participants

After signing the informed consent, 28 young adults (19-27, 23 ± 2, 12 male) and 21 older adults (60-75, 67 ± 4.70, 10 male) participated in the study. We excluded one older subject from analysis due to an error in task execution (wrong block order). All participants were right-handed as indicated by the Edinburgh inventory (Oldfield 1971) and were screened with general health and consumption habits questionnaires. Based on the general health questionnaire, participants with events, diseases or injuries that could affect the control of movement were excluded (e.g. head trauma). Based on the consumption habits, participants using recreational drugs or having hazardous alcohol consumption (more than 21 drinks per week for men or more than 14 for women) were excluded from the study.

No participants were excluded for these reasons. The older adults were assessed using the Mini-Mental State Examination and all scored within normal limits (score ≥ 24, (Folstein et al. 1975)). Approval was obtained by the Ethics Committee Research UZ / KU Leuven. Participants received financial compensation (€10/h).

Sample size was initially planned to reproduce the 20 participants per group as in Trewartha et al. 2014. We first included 20 older participants (recruited 21 but one was excluded due to error in the experiment) and 21 young ones. Upon data analysis, we noticed that, on average, younger people moved faster than older people in this paradigm despite the speed constraints (see below). We recruited an additional 7 young participants that were instructed to move slower to match hand velocity across age groups (as described below in the experimental paradigm).

### Experimental paradigm for the adaptation task

Participants made center-out, reaching movements in the horizontal plane while holding a robotic handle (KINARM End-Point Lab, BKIN Technologies). Hand position was represented by a white cursor on a display and vision of the hand was occluded. Movement trajectories were sampled at 1000 Hz. At the beginning of each trial, participants had to move their cursor to a starting point in the middle of the screen, after which a target appeared on one of eight possible locations (*Figure 1*) spaced ten cm away from the starting point. Participants were instructed to slice through the target by making a rapid, smooth reaching movement, avoiding any corrections. Once the movement amplitude exceeded ten cm, cursor position froze, providing feedback about movement accuracy and movement time. Movement times within 200 to 350ms resulted in a green cursor (for 5 young participants we increased the time window to 400 – 550ms and for another 2 to 300 – 400ms to keep average movement times the same for both age groups). Too slow movements caused the cursor to turn blue and too fast movements resulted in an orange cursor. After feedback, the starting point appeared, a new trial started, and the participants had to move their hand back to the starting position. On each trial, two points could be earned: one for hitting the target and one for applying the right speed. Participants were encouraged to collect as many points as possible.

**Figure 1:**
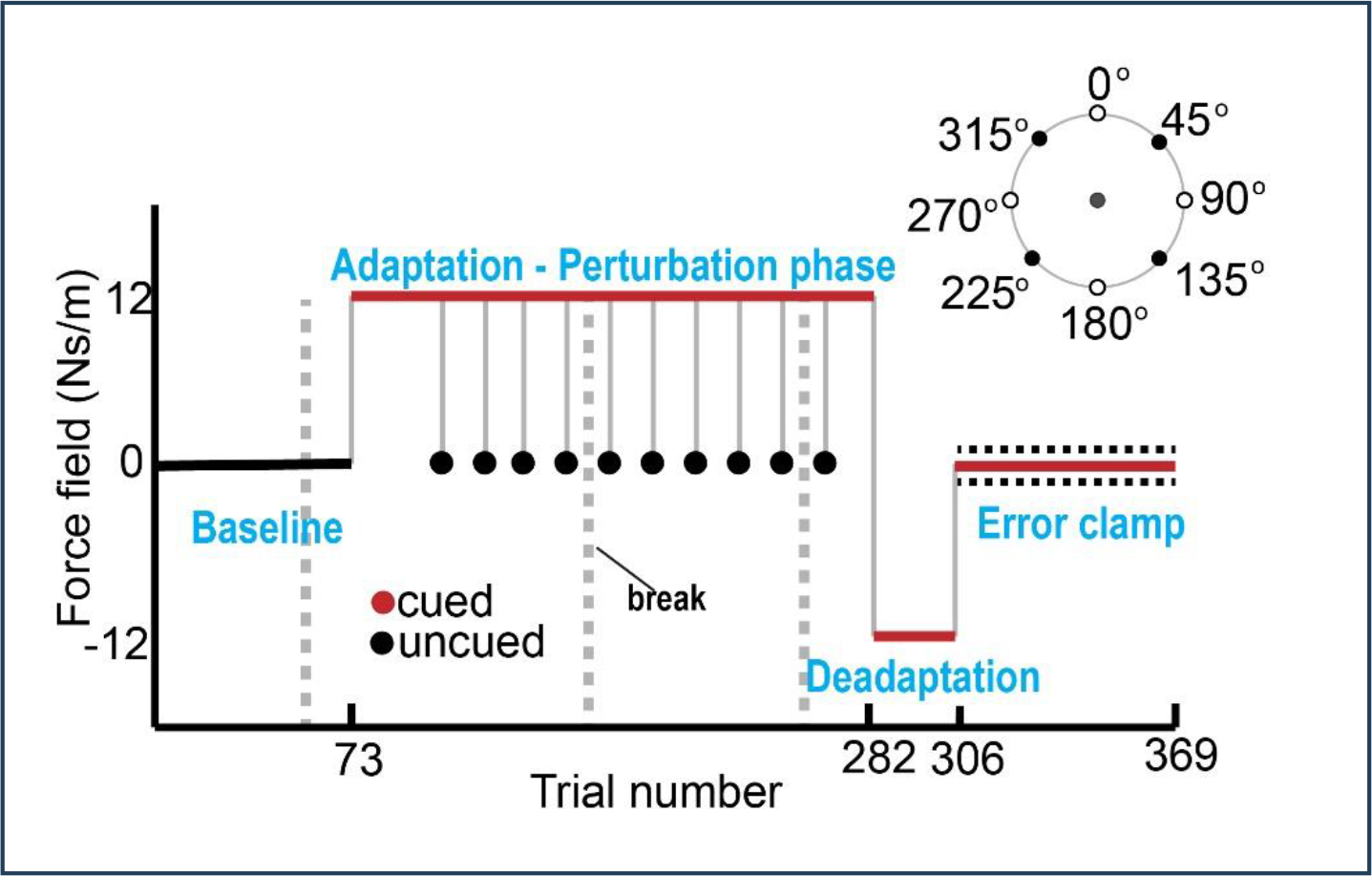
Paradigm. A change in cursor color indicated presence (cued) or absence (uncued) of a force field. Interspersed throughout the baseline and perturbation phase, were error clamp trials. Eight targets (open circles) were displayed, of which only four (filled black circles) were used for the uncued trials during the perturbation phase and error clamp trials during baseline and perturbation phase.

The task started with 72 baseline trials with reaches towards eight possible targets presented in pseudo-random order (9 cycles of 8 different targets) and a white cursor (Figure 1). Participants then continued with a perturbation phase (trials 73 – 281), during which a viscous force field (12 Ns/m) was applied perpendicular to hand velocity and the cursor had a red color (cued trials). Subjects received additional instructions, which were: “Initially, your cursor was a white dot, but from now on, your cursor can turn red. At that moment, something special will happen, but you still have to try to do the same thing, slice through the target with your cursor. A warning sign will be shown each time your cursor changes color”. Interspersed with these perturbation trials were trials with a white cursor (uncued trials), located on one of four possible locations. While the white cursor could be considered as a cue, we decided to adopt the terminology of Morehead and colleagues (2015). Participants could move straight ahead without interference of the force field and were occasionally reminded of this. From trial 282 to 305, the force field was reversed, washing out the adaptation to the first perturbation (Deadaptation in Fig.1). Lastly, retention was tested during an error-clamp phase (trials 306 – 369). Hand trajectory was constrained to a straight line from the starting point to the target, by guiding the handle between two stiff virtual walls (Scheidt et al. 2000). Visual feedback is provided during these trials. Throughout the baseline and perturbation phase, we pseudo-randomly introduced error-clamp trials to measure forces participants applied on the robot. While all 8 targets were used during the retention phase, only 4 of them were used for the error-clamp trials during the baseline and perturbation phase. The direction of the force field, clockwise or counterclockwise, was counterbalanced across subjects and three one-minute breaks were provided (dashed grey vertical lines on Fig.1, after trials 54, 153 and 253).

### Experimental paradigm for the visual working memory task

Given the importance of working memory for the explicit component of motor adaptation (Christou et al. 2016; Vandevoorde and Orban de Xivry 2020) and its potential link with the spontaneous recovery of motor memory (Trewartha et al. 2014), we decided to measure working memory capacity in all participants. It was quantified with a computer-based task (Christou et al. 2016; Saenen et al. 2022; Vandevoorde and Orban de Xivry 2019, 2020). Sixteen white lined squares were presented in a circular array with, in the middle, a white fixation cross. Three to six red circles were presented for two seconds randomly each in one of the squares. The array disappeared leaving only the fixation cross for three seconds, where after the squares returned with a question mark placed in one of them. Subjects had to indicate whether the probed location had contained a red circle. After three seconds of response time, a new trial began. Participants completed 48 trials (12 trials/condition) after eight practice trials. Two participants did not perform this task.

### Data processing and analysis

The x and y positions of the handle and x and y forces exerted on the handle were recorded at 1000 Hz. To combine the data from subjects who started with a clockwise force field with those who started with a counterclockwise force field, all the signs of position and force data in de x-direction for the clockwise condition were flipped.

For each field trial, lateral deviation from the optimal trajectory from starting point to target was calculated at peak velocity. Total adaptation level was quantified as the lateral deviation of the last 80 cued trials during the perturbation phase. Implicit learning was quantified as the average of the lateral deviation over the last 12 uncued trials during that phase.

In the error-clamp trials, the force subjects exerted on the channel walls (perpendicular to the heading direction) at peak velocity was used as a measure of adaptation. A second method we used to quantify learning in error-clamp trials was to compute the slope of the relationship between ideal force during reaching and the exerted lateral force (Smith et al. 2006; Trewartha et al. 2014). All trials from the perturbation phase onward were corrected for baseline error per target location. Retention of implicit learning, spontaneous recovery, was calculated as the average of the last 48 trials during the error-clamp phase. These outcome measures were controlled for peak hand velocity, because movement speed influences adaptation level (Shadmehr and Mussa-Ivaldi 1994). In addition, the slope of the relationship between ideal force and actual generated force was used as a control measure for the level of spontaneous recovery (adaptation index, Trewartha et al. 2014).

Working memory capacity is calculated using the following formula: K = S ^*^ (H – F). K is the memory capacity, S is the size of the array, H is the observed hit rate and F is the false alarm rate (Vogel et al. 2005). To estimate K, we used the decision tree used by Vandevoorde et al. (2020) and published as supplementary material by Saenen et al. (2022) and available at https://doi.org/10.6084/m9.figshare.23535396.v1.

For statistical testing, we used t-tests with unequal variance in all tests. All statistical tests were also reproduced with non-parametric tests but the results between the parametric and non-parametric tests never differed in their conclusion. Effect sizes (Robust Cohen’s d) and its confidence interval (computed with bootstrap with 5000 iterations) were obtained from the meanEffectSize function in Matlab. ANCOVA’s were performed with the aoctool function in matlab (with the model ‘parallel lines’), fitting a separate line to each group, but constraining these lines to be parallel as we did not expect a different relationship between the covariates and the dependent variables in function of age. For all the analyses, the α-level was set at 0.05.

To test for the absence of age-related difference in adaptation (hypothesis 1), we compared the average lateral deviation at the end of the adaptation period over the last 80 cued field trials (analysis 1), the average exerted force (measured at peak velocity) during the last 12 cued clamped trials of the adaptation phase (analysis 2) and the implicit adaptation level measured as the average lateral deviation over the last 12 uncued trials (analysis 3) between young and older participants with an independent t-test with unequal variance. Additionally, an ANCOVA was used to check for any influence of hand velocity on these outcomes. The outcome was set as dependent factor and hand velocity for these specific trials were used as covariate. For each of the analyses, we also performed a Bayesian independent T-test with a Cauchy distribution as prior (width of 0.707).

In analysis 4, we tested age-related differences in spontaneous recovery level measured either as the average force exerted at peak velocity or as the average adaptation index computed over the last 48 clamped trials of the error-clamp period. These two outcomes were submitted to the same statistical tests as in analysis 1. The force data was also submitted to a Bayesian independent Samples T-test in JASP to test how compatible this data was with the hypothesis that spontaneous recovery was larger in young than in older participants (one-sided t-test). The selected prior for this analysis was the default Cauchy prior (Scale=0.707). The Bayesian analysis was performed in JASP (JASP Team 2023).

To test possible correlation between adaptation levels at the end of the perturbation phase and during the error-clamp period (hypothesis 3), implicit adaptation levels at the end of the perturbation period (from analysis 3) and spontaneous recovery levels (from analysis 4) were correlated via multilevel correlation (Analysis 5) from the correlation package in R (Makowski et al. 2020). The different levels corresponded to the different age groups.

In analysis 6, an additional ANCOVA was used with spontaneous recovery level (from analysis 4) as dependent factor and implicit adaptation level at the end of learning (from analysis 3) as covariates.

In analysis 7, we conducted a Bayesian independent t-test in JASP (JASP Team 2023). The prior was centered on the effect size reported in the original study by Trewartha et al (d=0.8) and followed a Cauchy distribution. We used three different scales for the prior in order to test the sensitivity of our results to the width of the prior distribution. In this analysis, we test the hypothesis that the difference in spontaneous recovery level between age groups is equal to d=0.8.

In analysis 8, working memory capacity was compared between young and older participants with an independent t-test with unequal variance.

In analysis 9, we investigate the possible association (hypothesis 4) between working memory capacity (from analysis 8) and spontaneous recovery levels (from analysis 4) via multilevel correlation from the correlation package in R (Makowski et al. 2020). The different levels corresponded to the different age groups.

All data can be found on the RDR repository of the KU Leuven: https://doi.org/10.48804/KMGKLH All analysis scripts can be found at: <u>10.5281/zenodo.8284036</u>

## Results

### Force-field adaptation does not decline with aging

The aim of this experiment was to measure the impact of aging on implicit adaptation and its short-term retention through spontaneous recovery. Participants made center-out reaching movements towards targets, while adapting to a force field that pushed their hand away perpendicular to the heading direction (cued trials with red cursor). With practice, subjects gradually decreased their error over the course of learning (*Figure 2A*). Total adaptation level at the end of the perturbation phase was similar across age groups (*Figure 2B*, mean ± SD, young: 2.23±1.43mm, older: 2.29±1.87mm, Analysis 1: t(34.07)= −0.13, p=0.89, Cohen’s d = −0.011 CI=[−0.64,0.62]) and at the end of the deadaptation period (trials 298 to 305, t(45.25)=0.75, p=0.46, Cohen’s d=0.378 CI=[-0.19, 1.02]).

**Figure 2:**
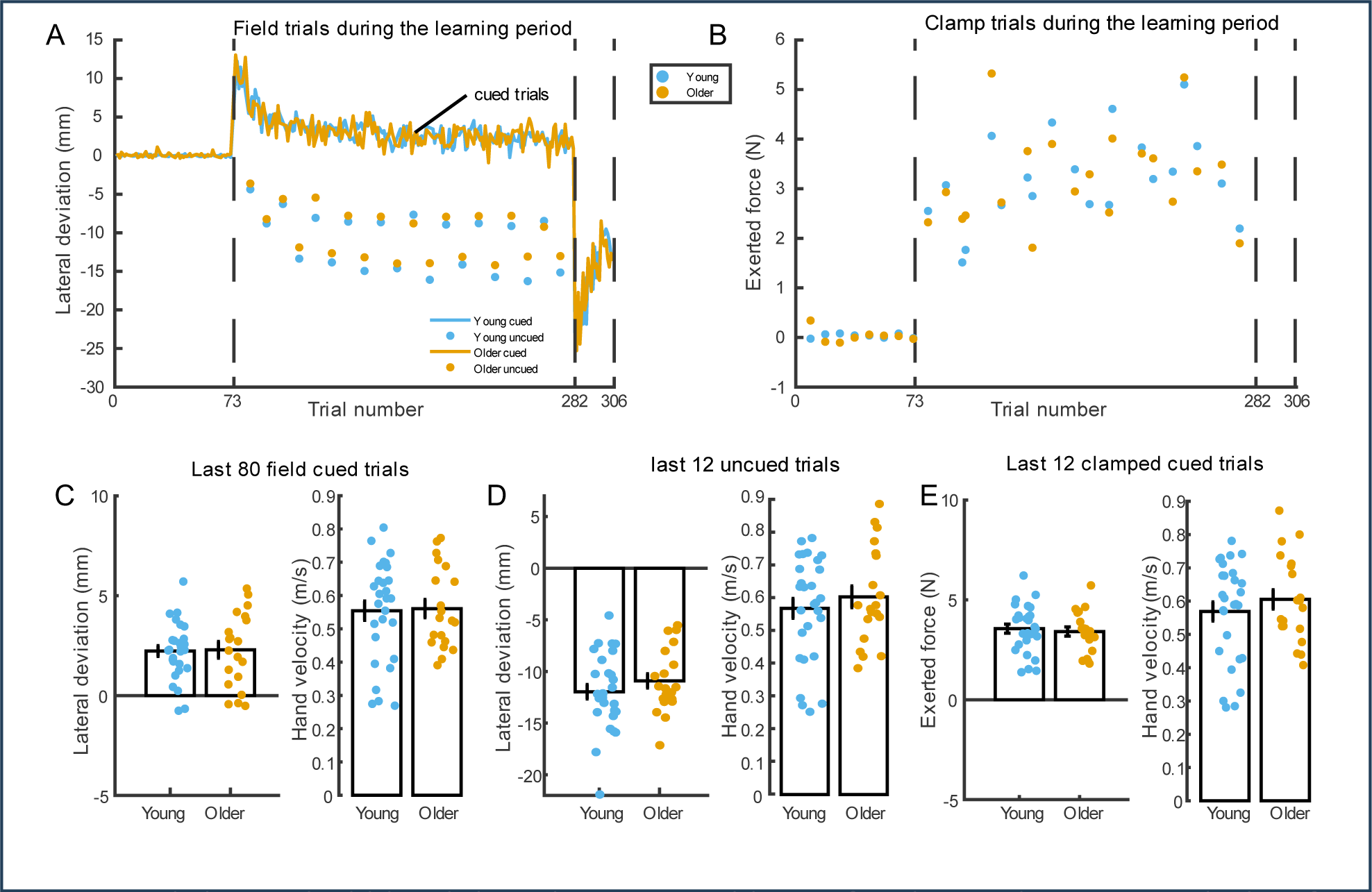
Total adaptation level did not decline with aging. **A** Lateral deviation from the optimal trajectory at peak velocity for young (blue) and older adults (orange) over the course of baseline and perturbation phase. Interspersed with these perturbation trials were uncued baseline trials (filled circles) where no perturbation was applied. **B** Exerted force perpendicular to heading direction at peak velocity during baseline and perturbation phase for young (blue) and older adults (orange). **C** Lateral deviation during the last 80 field trials from the perturbation phase and the corresponding hand velocity. **D** Lateral deviation during the last 12 uncued trials from the perturbation phase and the corresponding hand velocity. **E**. Exerted force during the last 12 error clamp trials from the perturbation phase and the corresponding hand velocity. For panels C, D and E, each dot represents the mean data from one individual. Error bar represents mean and standard error. For all panels, data from 28 young and 20 older participants are presented.

Given the importance of hand speed in force-field adaptation, we checked that the hand speed was comparable across the two groups. At the end of the adaptation period, hand velocity was comparable (Fig.1C, young: 0.55±0.15 m/s, older: 0.56±0.12 m/s, t(45.35)= −0.156, p=0.87, d = 0.053, CI=[−0.46,0.76]). Controlling movement speed did not change the outcome of the analysis of the lateral deviation at the end of the adaptation period (ANCOVA, F(1,45)=0.0161, p=0.899). The corresponding Bayesian analysis suggested that there was moderate evidence an absence of difference (BF = 0.293).

The force that participants exerted against the perturbation built up as participants learned to counteract the perturbation (*Figure 2B*). In error-clamp trials, the exerted force reached similar levels at the end of the perturbation phase for both groups (*Figure 2D*, mean ± SD, young: 3.6±1.2N, older: 3.4±1N, Analysis 2: t(44.15)= 0.46, p=0.65, Cohen’s d = 0.16, CI=[−0.42,0.79]). For those trials, we also did not find any evidence that the velocity varied across age groups (t(44.34)= −0.89, p=0.38, Cohen’s d = −0.104, CI=[−0.60,0.52]). Therefore, controlling for hand speed did not change the results (Analysis 2, ANCOVA: F(1,45)=1.84, p=0.18). The corresponding Bayesian analysis suggested that there was moderate evidence an absence of difference (BF = 0.315).

In some trials (uncued trials), we warned the participants that the force field would be turned off in order to forced participants to stop using any strategy to compensate for the perturbation and to measure implicit adaptation (Morehead et al. 2015). In these trials, perpendicular error increased with continued learning (*Figure 2A*). Participants made reaching movements to four different target locations (ordinal directions, see figure 1). These trials were randomly presented throughout the perturbation phase, but in a fixed sequence. For some reason, participants from both groups exhibited different amount of lateral deviations in function of target direction (Figure 2A). However, this effect of target direction was identical between the two age groups. We averaged the responses of the last 12 uncued trials and compared these between our age groups (*Figure 2E, Analysis 3*). We could not find evidence for a difference in implicit adaptation between young and older adults (mean±sd, young:-11.96±3.70mm, older: −10.91±3.09mm, Analysis 3: t(44.76)= −1.07, p=0.29, Cohen’s d = −0.24, CI=[−0.83,0.33]). In those trials, we did not find any evidence that hand speed differed across groups (young: 0.573±0.162 m/s, older: 0.609±0.149 m/s, t(43.00)= −0.785, p=0.4368, Cohen’s d = −0.076, CI=[−0.56,0.61]). This result remained the same after the implicit adaptation level was controlled for movement speed (Analysis 3: ANCOVA: F(1,45)=2.18, p=0.146). The corresponding Bayesian analysis suggested that there was anecdotal evidence an absence of difference (BF =0.45).

### No evidence that spontaneous recovery declines with aging

At the end of the experiment, lateral deviation of each movement was clamped to zero, ensuring participants would always hit the target. This enabled us to measure the retention of implicit adaptation without interference of trial-by-trial learning. Exerted force increased over time in the same direction as during the perturbation phase, characteristic of spontaneous recovery (*Figure 3A*). The average response of the last 48 clamp trials were compared between age groups (*Figure 3B*) and we did not find any evidence for a difference between young and older adults (median, young: 0.88±0.73N, older: 1.01±1.5N, Analysis 4: t(25.46)= −0.35, p=0.73, Cohen’s d = 0.036, CI=[−0.54,0.67]). Note that this result is independent of which trials are analyzed. Performing a trial-by-trial analysis as in Trewartha et al, no between-group differences remained significant after correction for multiple comparisons (p<0.05/64). Similarly, analyzing all 64 trials from the spontaneous recovery period provided the same statistical results (t(25.44)= −0.367, p=0.72, Cohen’s d = 0.003 CI=[−0.58,0.71]). We did not find any evidence that hand speed differed across groups (young: 0.53±0.14 m/s, older: 0.55±0.09 m/s, t(45.99)= −0.4, p=0.69, Cohen’s d= 0.034, CI=[−0.46,0.78]), indicating we succeeded in this aim.

**Figure 3:**
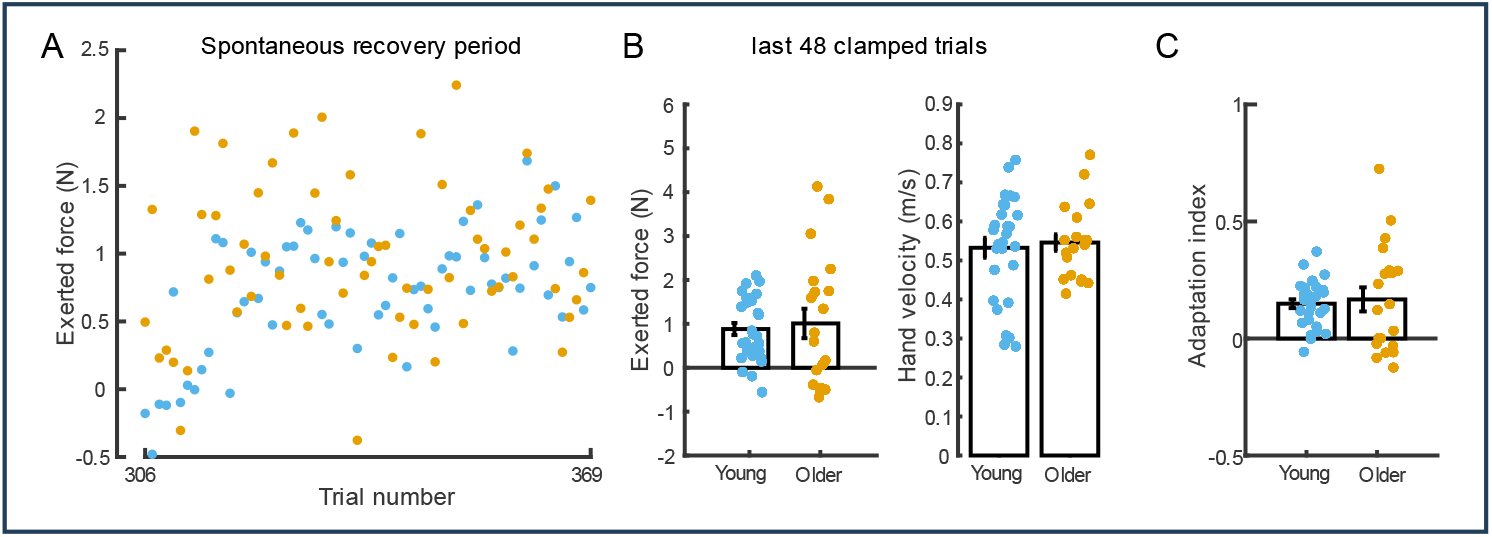
Spontaneous recovery did not decline with aging. **A** Exerted force at peak velocity during the error-clamp phase for young (blue) and older adults (orange). Each dot represents the average force exerted by the individuals from an age group for a single trial. **B** Exerted force during the last 48 trials of the error-clamp phase. **C** Adaptation index for the last 48 trials of the error-clamp phase. For panels B and C, each dot represents the mean data from one individual. Error bar represents mean and standard error. For all panels, data from 28 young and 20 older participants are presented.

Controlling movement speed did not change the result (Analysis 4: F(1,45)=0.138, p=0.71). In addition, the adaptation index (Figure 3C), which was used in Trewartha et al. (2014), did not differ between age groups either (young: 0.15 ± 0.1, older: 0.17 ± 0.23, Analysis 4: t(24.20)= −0.335, p=0.7402, Cohen’s d = 0.014, CI=[−0.56,0.65]).

To confirm these results, we performed a Bayesian analysis on the force data in order to test how compatible our data was with the idea that the spontaneous recovery level was larger in young than in older participants. There was moderate support (BF=4.44) for the idea that the spontaneous recovery level was not larger in young than in older participants.

This failure to replicate the effect described in Trewartha et al. is consistent with the fact that we did not find any evidence for a difference in implicit adaptation between the two age groups in this study (Fig.2) and in previous studies (Vandevoorde and Orban de Xivry 2019, 2021) if the level of spontaneous recovery is linked to the level of adaptation at the end of the learning period. Indeed, we expect that people with more implicit adaptation at the end of learning exhibit more spontaneous recovery, resulting in a positive correlation between the two. Therefore, we pooled the data for all participants and correlated both measures while taking the two different groups into account (*Figure 4, Analysis 5*). A significant positive correlation was found between the level of implicit adaptation and the level of spontaneous recovery (N = 48, r = 0.55, t(46)=4.42, p < 0.001).

**Figure 4:**
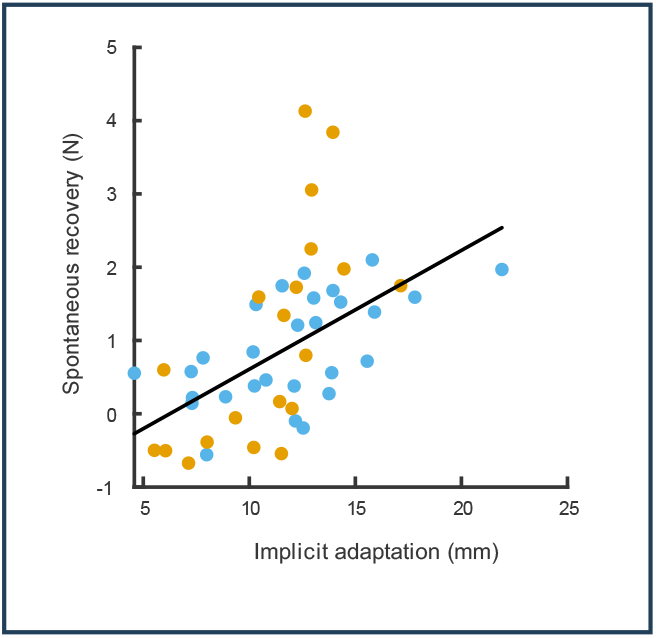
Implicit adaptation (data from figure 2.D) and spontaneous recovery (data from figure 3.B) are correlated (N = 48). To facilitate interpretation, implicit level was converted to positive values, such that participants with a higher implicit adaptation level have a larger lateral deviation. Each dot represents the data from one individual. Data from both groups were combined thanks to multilevel correlation. Regression line was obtained with robustfit method in Matlab.

Given this correlation, it might be that small differences in implicit adaptation level at the end of the learning period can mask age-related effects in spontaneous recovery. That is, if older participants had slightly larger implicit adaptation levels, it could compensate for a decrease in spontaneous recovery. Therefore, we compared spontaneous recovery across age groups while controlling for implicit adaptation levels (Analysis 6). Yet, this additional analysis further confirmed our previous result and did not provide any evidence that spontaneous recovery level was smaller in older participants (F(1,45)=1.317, p=0.2571). If anything, marginal means obtained in the ANCOVA tended to indicate that, when controlling for implicit adaptation levels, older adults tended to exhibit more spontaneous recovery than younger adults (young: 0.88N±0.73; older: 1.009±1.5, mean±SD).

### Combining the data of the original study and of the present conceptual replication favor the null hypothesis

The effect size for the difference in force used between young and older subjects during spontaneous recovery in the study of Trewartha was d=0.8 (personal communication from Trewartha). We use this effect size as a prior with Cauchy distribution. In this case, a Bayes Factor (BF_10_) larger than 1 would favor the effect size found in the original study (favoring the hypothesis that the difference in spontaneous recovery between young and old participants is as big as claimed by Trewartha et al. (d=0.8)). A Bayes Factor smaller than 1 would indicate that the effect is smaller than in the original study (favoring H0). As a sensitivity analysis, we tested different widths for the prior distribution (narrow 0.5, medium: 0.707, wide: 1). In all cases, the posterior was closer to 0 than the prior, with Bayes Factor (BF_10_) yielding substantial evidence (BF10>3, Dienes 2014) that the difference in spontaneous recovery levels should be smaller than d=0.8 (BF_01_ = 6.89 for a default prior width = 0.707, BF_01_ = 7.33, narrow prior with SD = 0.5; BF_01_ = 7.3, wide prior with SD = 1.414). Overall, the hypothesis that the effect of age on spontaneous recovery level is smaller than 0.8 was 6 to 7 times more likely than an effect size of 0.8. In other words, the Bayesian analysis favored the hypothesis that the actual effect of aging on spontaneous recovery was smaller than that of the original study with a median effect size of 0 and a confidence interval of [-0.58, 0.56].

### No correlation between explicit adaptation level and working memory capacity score

In the study of Trewartha et al. (2014), spontaneous recovery level were linked to cognitive processes such as explicit memory (their Fig.7). Therefore, we checked whether we could link any aspects of spontaneous recovery to explicit memory processes such as working memory capacity. In our sample, we tested working memory capacity in all our participants except two young adults. Older adults exhibited lower working memory capacity than younger adults (Analysis 8, t(42.26)= 3.5, p=0.0011, Cohen’s d = 1.16, CI=[ 0.48,2]). Yet, this does not seem to affect the amount of spontaneous recovery as this was similar across age groups (Fig.3). In addition, we did not find any evidence that the amount of spontaneous recovery was correlated with the working memory capacity score (Analysis 9, Fig. 6, N = 46, r = −0.05, t(44)=-0.36, p = 0.72). This questions the link between memory processes and spontaneous recovery of motor adaptation.

**Figure 5:**
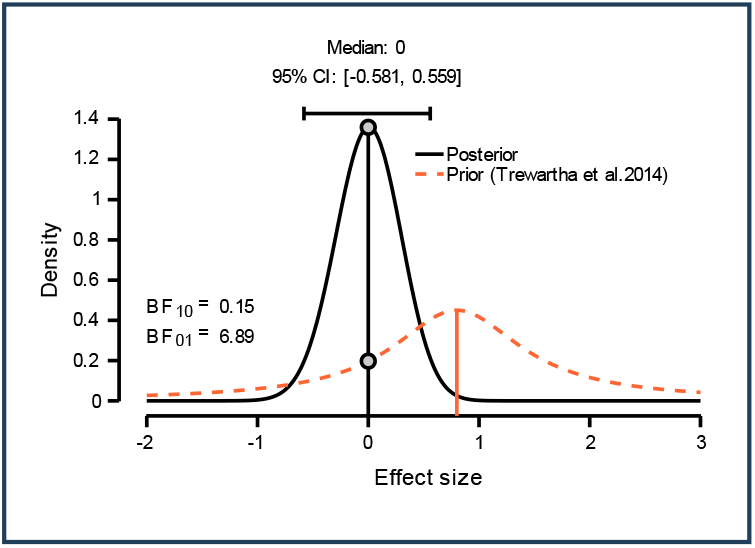
Output of the Bayesian analysis. The Bayesian analysis takes the previous data as the prior (centered on d=0.8, Trewartha et al. 2014) and computes the posterior based on the data of the present study.

**Figure 6:**
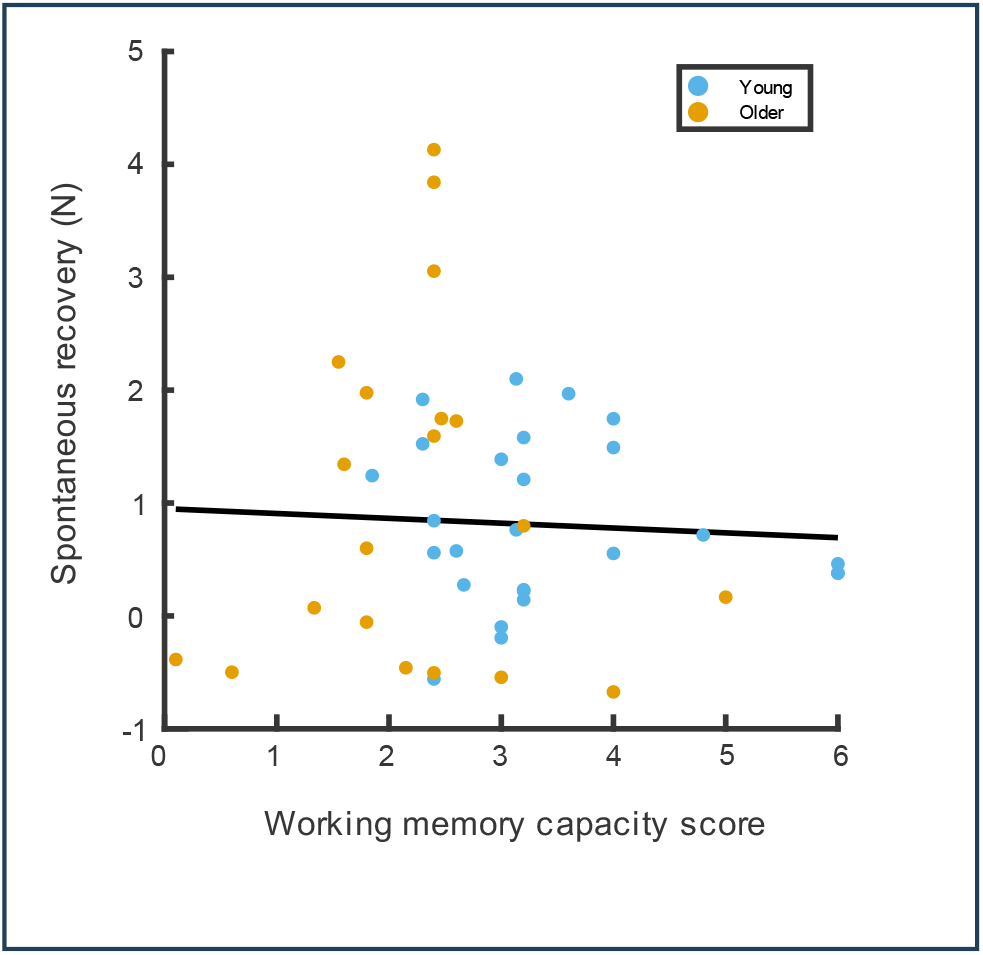
Spontaneous recovery level was not correlated with spatial working memory capacity score (N = 46). Each dot represents the data from one individual. Data from both groups were combined thanks to multilevel correlation. Regression line was obtained with robustfit method in Matlab.

## Discussion

In this study, we tested whether aging influenced the ability to adapt reaching movements accordingly when movements were perturbed. Participants reached to targets while a force field perturbed their movements in an adaptation period. In some catch trials, participants were cued that the force field would be turned off in the subsequent trial (Morehead et al. 2015). Any error in reaching direction in these trials was attributed to implicit adaptation. After a short de-adaptation period with a reversed force field, spontaneous recovery of motor memories of the adaptation period was tested by guiding the hand directly towards the targets in error clamp trials (Smith et al. 2006). Across age groups, we observed little difference in performance in this task. Both total adaptation and implicit adaptation were not impaired in older adults compared to their younger controls. In addition, we failed to replicate the observation of Trewartha and colleagues (2014) and found that spontaneous recovery remained also unaffected by aging. Yet, implicit adaptation and spontaneous recovery levels were correlated independently of age groups, suggesting that spontaneous recovery is linked to the memory of implicit adaptation (McDougle et al. 2015). In contrast to Trewartha et al. (2014), we could not find any evidence that spontaneous recovery of motor memories were linked to memory processes.

Our results potentially resolve the contradiction that spontaneous recovery, but not implicit adaptation, was impaired with aging (Trewartha et al. 2014; Vandevoorde and Orban de Xivry 2019). Indeed, the slow process of adaptation is believed to reflect the implicit process (Mazzoni and Krakauer 2006; Morehead et al. 2017) and the spontaneous recovery is linked to this slow process (McDougle et al. 2015). It was therefore surprising that some studies found that the implicit component of adaptation was not affected by aging (Heuer and Hegele 2008; Huang et al. 2017; Vandevoorde and Orban de Xivry 2019) but that the spontaneous recovery was (Trewartha et al. 2014), given that they come from the same process (McDougle et al. 2015; Smith et al. 2006).

The absence of age-related impairment in spontaneous recovery implies that as we age, we do not get more forgetful of movements in the short-term. Indeed, spontaneous recovery is a measure of short-term retention of the slow implicit process. Following the two-state model (Smith et al. 2006), the spontaneous recovery results from the rapid decay of the fast state to zero in the error-clamp phase, while the slow process still contains a memory trace of the motor memory acquired during the first adaptation phase. This is consistent with the correlation between the amount of implicit adaptation during learning and the amount of spontaneous recovery (Fig. 5). This is also consistent with the findings of McDougle et al (2015). Therefore, a decrease in spontaneous recovery could be due either to worse implicit adaptation during learning (which we did not find) or smaller retention rate (Bindra et al. 2021). The absence of age-related difference in spontaneous recovery suggests that there is no evidence for an age-related deficits in either implicit adaptation or retention rate. This finding is in contrast to the results reported by Trewartha et al (2014) who found lower spontaneous recovery in older people.

Previous studies that quantified short-term retention in old and young adults gave mixed results. No deficit in short-term retention measured after a one-minute break was reported in a visuomotor rotation task (Vandevoorde and Orban de Xivry 2019). These authors investigated retention of visuomotor adaptation in two different adaptation paradigms. First, one-minute breaks were inserted during regular visuomotor rotation paradigm. In this case, there was no evidence of a difference in retention level of total adaptation between young and old participants. Second, they used one-minute breaks during task-irrelevant clamped feedback paradigm that is known to elicit pure implicit adaptation (Avraham et al. 2021; Kim et al. 2019; Morehead et al. 2017; Morehead and Orban de Xivry 2021). In this case again, there was no evidence for a deficit in short-term retention of implicit adaptation. However, one other study that measured the explicit component of visuomotor rotation by asking participants to report their aiming direction found that older participants exhibited worse retention of implicit adaptation (Bindra et al. 2021). Yet, it is unclear why people would change the explicit report of their aiming direction in a one-target task after a one-minute break if nothing happened during the break. Similarly, an age-related deficit in the retention did occur in a gait adaptation paradigm (Malone and Bastian 2015). These authors suggested that the implicit, and not the explicit component of adaptation was impaired, because larger forgetting was observed in older adults independently of whether a cognitive distraction was presented during the gait adaptation period or not. Such a cognitive distractor would have the ability to reduce the contribution of the explicit component. For this reason, the observed effect was indirectly attributed to the implicit component of adaptation.

### Possible sources of discrepancy with the study of Trewartha

The fact that our results differ from the study of Trewartha et al. (Trewartha et al. 2014) might stem from one of the small differences in protocol between our studies even though we tried to use a very similar protocol to theirs. Yet, they differed in several aspects.

The experimental design of the forcefield task here used 8 radial targets from a central start position. In contrast, the Trewartha study used alternating movements between two targets, with forces only applied to movements in one direction. The impact of target number on age-related differences in implicit motor adaptation (or absence thereof) remains unknown. Our protocol had a more extensive adaptation period (209 trials vs. 118 in Trewartha et al.), deadaptation phase (24 vs. 15) and retention phase (63 vs. 22), while the baseline and de-adaptation phases were similar (baseline 73 vs. 52 trials and de-adaptation 24 vs 20). The longer adaptation period might have resulted in more opportunity for the participants to learn the force field implicitly, which might have concealed a learning deficit in the older adults that is then later reflected in the spontaneous recovery period. However, Trewartha et al. did not observe any difference in adaptation level during learning.

The type of movement and allowed movement speed also differed across the studies. While our participants had to slice through the target, those from Trewartha et al. had to stop on the target. We allowed for a greater variability in hand velocity, allowing faster movements (0.3 – 0.5 m/s vs 0.3-0.4 m/s in the Trewartha et al.). Yet, the hand velocity was matched between our group of young and old participants. Because older adults tended to move slower, we asked a few young participants to perform the experiment while adapting the accepted speed range. On average, our age groups moved with the exact same velocity.

One additional difference lies in our sample. Trewartha et al. showed that, within their sample, participants who scored high on an explicit memory task had better spontaneous recovery. So maybe, our sample of older people all had very good cognitive memories. Yet, our sample of older participants had worst working memory capacity than younger participants.

Finally, Trewartha measured explicit and implicit components in separate tasks and compared the results between age groups. Older adults scored less in both the explicit and implicit task. We measure implicit adaptation within our adaptation paradigm during learning and working memory in a separate task. This test of implicit adaptation could also have influenced the outcomes of the study. We found no difference in implicit level even though older adults had worse working memory capacity.

For all these reasons, our study represents a conceptual replication of the study by Trewartha et al. 2014 and not a direct/exact replication. If any of the factors identified as differences between our study and that of Trewartha is responsible for the difference in outcomes, it means that the age-related effect on spontaneous recovery, if it exists, is highly sensitive to the experimental conditions. By employing multiple methodologies, conceptual replications provide a robustness test of the findings. Our study suggests that the generalizability and robustness of the original results should be considered with caution. The age-related difference in spontaneous recovery found by Trewartha and colleagues might be true, but is likely dependent on the experimental conditions.

### Do explicit/cognition or implicit adaptation relate to spontaneous recovery

Beyond the technical differences, there are also differences in the theoretical approaches between the two studies. While Trewartha focused on the role of cognition on the spontaneous recovery, we believe that implicit motor adaptation modulates spontaneous recovery. Indeed, Trewartha and colleagues found that people who had “good” explicit memory had higher levels of spontaneous recovery. Our attempt at a conceptual replication of this correlation failed as we did not find any evidence that working memory capacity was linked to spontaneous recovery. The result of Trewartha and colleagues is at odds with the study of Keisler and colleagues (2010) who showed that a secondary cognitive task disrupted the fast process but not the slow process responsible for spontaneous recovery (McDougle et al. 2015). Our results rather agree with the results of Keisler than with those of Trewartha. Indeed, we found that the amount of implicit adaptation measured during learning correlated with the level of spontaneous recovery across participants.

### Statistical view on this absence of replication

Our study and the study of Trewartha provide conflicting results. Our Bayesian analyses aimed at reconciling those conflicting results. The Bayesian analysis suggests that, given our data, the influence of age on the spontaneous recovery of motor memories is very likely much smaller than what was reported by Trewartha and colleagues (2014). Yet, the Bayesian analysis does not prove that there is no effect. It estimates that the effect size lies somewhere in an interval between −0.6 (medium effect size of larger spontaneous recovery for older people) and 0.55 (medium effect size for a larger spontaneous recovery for younger people.

Yet, beyond such statistical arguments, our results are well aligned with the observation that the spontaneous recovery of motor memories depends on the slow implicit component of motor adaptation (Keisler and Shadmehr 2010; McDougle et al. 2015; Smith et al. 2006) and that this component is not affected by aging (Cressman et al. 2010; Hegele and Heuer 2010, 2013; Heuer and Hegele 2008; Huang and Ahmed 2014; Kitchen and Miall 2021; Reuter et al. 2020; Vandevoorde and Orban de Xivry 2019, 2021). The results of Trewartha and colleagues are at odds with this theory, which motivated our conceptual replication attempt.

### Limitations of the study

In this study, we measured the implicit component of motor adaptation by looking at the distance participants deviated from the straight trajectory. However, short-term retention was measured by the force that was exerted perpendicular to the heading direction. This difference in units makes direct comparison between the two measures difficult. Another way of separating implicit from explicit learning is described by Sween et al. (2020). Implicit adaptation level was determined with ‘No Push’-trials, where participants were instructed to ignore the force field and to not push against it. Total adaptation, including the explicit component, was measured in ‘Push’-trials, which had an extra reminder to push against the force field. The difference in exerted force is attributed to the explicit component of motor adaptation. The results indicate that in a ‘Push’-trial, participants apply more force and in a ‘NoPush’-trial less force, as compared to a regular trial. Therefore, our study could have benefited from such an assessment.

Our Bayesian analysis suggests that the maximum effect size should be much smaller than anticipated based on the study of Trewartha. This means that we only have 60% power to detect an effect if there is one of d=0.6. This should motivate future studies to include more participants as we now have a better estimate of the possible effect size range.

Finally, the group of older participants exhibited much more inter-subject variability than the group of younger participants. This is typical in aging studies but would need to be tackled to get a better estimate of the spontaneous recovery of these older participants.

## Conclusion

We attempted a conceptual replication of the effect of age on spontaneous recovery as demonstrated by Trewartha and colleagues but could not replicate their results as we failed to find evidence for a difference in spontaneous recovery between young and old participants. The current results are more in line with the idea that spontaneous recovery depends on the retention of implicit adaptation and that implicit adaptation is not affected by aging.

## Acknowledgement

We thank Yee Kee Lam for help with data collection. We thank Matheus Pacheco for comments on an earlier version of the manuscript. The authors declare no competing interests. A preprint version of this article has been peer-reviewed and recommended by PCI Health & Mov Sci (https://doi.org/10.24072/pci.healthmovsci.100040).

